# Multiplexed VaxArray Immunoassay for Rapid Antigen Quantification in Measles and Rubella Vaccine Manufacturing

**DOI:** 10.1101/2021.03.05.433809

**Authors:** Jacob H. Gillis, Keely N. Thomas, Senthilkumar Manoharan, Mallikarjuna Panchakshari, Amber W. Taylor, David F. Miller, Rose T. Byrne-Nash, Christine Riley, Kathy L. Rowlen, Erica Dawson

**Affiliations:** InDevR Inc., Boulder, Colorado, USA; Biological E. Ltd., Hyderabad, India; GT Molecular, Fort Collins, CO

**Keywords:** Measles, Rubella, CCID_50_, Identification, Antigen Quantification, MR Vaccine, Vaccine Manufac turing, Multiplexed Immunoassay, Microarray, VaxArray

## Abstract

Measles-containing vaccines (MCV), specifically vaccines against measles and rubella (MR), are extremely effective and critical for the eradication of measles and rubella diseases. In developed countries, vaccination rates are high and vaccines are readily available, but continued high prevalence of both diseases in developing countries and surges in measles deaths in recent years have highlighted the need to expand vaccination efforts. To meet demand for additional vaccines at a globally affordable price, it is highly desirable to streamline vaccine production thereby reducing cost and speeding up time to delivery. MR vaccine characterization currently relies on the 50% cell culture infectious dose (CCID_50_) assay, an endpoint assay with low reproducibility that requires 10-14 days to complete. For streamlining bioprocess analysis and improving measurement precision relative to CCID_50_, we developed the VaxArray Measles and Rubella assay kit, which is based on a multiplexed microarray immunoassay with a 5-hour time to result. Here we demonstrate vaccine-relevant sensitivity ranging from 345 – 800 IFU/mL up to 100,000 IFU/mL and specificity that allows simultaneous analysis in bivalent vaccine samples. The assay is sensitive to antigen stability and has minimal interference from common vaccine additives. The assay exhibits high reproducibility and repeatability, with 15% CV, much lower than the typical 0.3 log_10_ error (~65%) observed for the CCID_50_ assay. The intact protein concentration measured by VaxArray is reasonably correlated to, but not equivalent to, CCID_50_ infectivity measurements for harvest samples. However, the measured protein concentration exhibits equivalency to CCID_50_ for more purified samples, including concentrated virus pools and monovalent bulks, making the assay a useful new tool for same-day analysis of vaccine samples for bioprocess development, optimization, and monitoring.

## INTRODUCTION

Measles and rubella are highly contagious viruses that cause significant morbidity and mortality. Measles virus is one of the most infectious human-borne viruses and, prior to widespread vaccination, infected > 90% of children before 15 years of age [1]. Acute measles infection is associated with fever, cough, and rash that can persist for a week [2]. Up to 6% of acute measles cases are fatal and the World Health Organization (WHO) estimates 535,000 children, most in the developing world, died from acute measles infection in 2000 [3]. Subacute sclerosing panencephalitis (SSPE), a chronic complication of measles infection, can occur in patients up to 10 years after initial acute infection. While rare (0.007-0.011% of infections [4]), SSPE is always fatal, usually within 1 to 3 years.

Unlike measles, acute rubella infection is rarely fatal. Rubella infection during the first trimester of pregnancy is associated with severe chronic developmental disorders (congenital rubella syndrome, CRS) in the developing fetus that includes cataracts, deafness, encephalopathy, heart defects, and severe mental development disorders. When infected during the first 12 weeks of gestation, up to 80% of fetuses will develop CRS [5,6]. Prior to introduction of rubella vaccination (and in non-immunized populations), CRS occurs at a rate of 0.8-4.0 per 1000 live births [7], though this likely greatly under-reports the occurrence [8].

Current measles vaccines are up to 99% effective, but high coverage rates of 95% or greater are necessary for elimination of measles [9–11]. Rubella vaccines have demonstrated similar efficacy [8,12,13], reducing the incidence of CRS in newborns to < 20 total per year in WHO regions with high (>90%) vaccination coverage [14].

Measles-containing vaccines (MCV) are affordable, available, and have been in use for a half-century in the developed world, but coverage in low- and middle-income countries is lacking due in part to cold-chain requirements and manufacturing costs. Several initiatives aim to enable low-cost manufacturing of measles and rubella (MR) vaccines at the quantity and cost required for widespread vaccination in the developing world [15]. Recent efforts have been highly successful, reducing the number of measles-associated deaths by 74% worldwide between 2000 and 2010 [16], but a 50% increase in measles mortality worldwide between 2016 and 2019 [16] highlights the fight is not over.

MR vaccines rely on cell-culture based assays, such as 50% Cell Culture Infectious Dose (CCID_50_), for characterization during manufacturing, including identity, potency, and stability testing. Infectivity measurements are widely adopted and highly predictive of vaccine efficacy [17–20] but present certain limitations, as the time to result is 10-14 days and execution requires trained personnel. In addition, CCID_50_ assays rely on subjective, discontinuous endpoint measurements and are notoriously error prone with observed variability of ± 0.3 log_10_ infectious units per mL (50-100% variability) [21–23], resulting in costly hold times and lot rejections that can increase manufacturing costs and delay delivery of critical vaccine doses.

This work describes the development of an alternative assay for tracking antigen content, stability, and identity throughout the MR vaccine manufacturing process. We have adapted the VaxArray technology, described previously for influenza vaccines [24–27], for the evaluation of MR vaccine antigens. VaxArray assays are simple multiplexed immunoassays that use monoclonal antibodies in a glass microarray format. In addition to multiplexing, VaxArray provides a significant advantage in terms of reagent use, requiring 100x less capture antibody than ELISA. Here we demonstrate feasibility of the VaxArray technology for measuring antigen content in MR vaccines, enabling streamlined characterization at several manufacturing steps.

## MATERIALS AND METHODS

### Anti-measles and rubella antibodies

Antibodies raised against measles and rubella virus-specific were obtained from numerous commercial sources, including Lifespan Bioscience (Seattle, WA), Novus Biologicals (Littleton, CO), Fitzgerald Industries International (Acton, MA), GeneTex (Irvine, CA), ViroStat Inc. (Westbrook, ME), Antibodies-Online (Aachen, Germany), MilliporeSigma (Burlington, MA), and HyTest Ltd (Turku, Finland).

### VaxArray MR assay

The VaxArray Measles and Rubella (MR) assay is similar to previously-described VaxArray Influenza assays [24–27], with the slide layout, microarray layout, and detection principle depicted in **Figures 1a, b**, and **C**, respectively. Virus-specific monoclonal antibodies are printed on the microarray (**Figure 1A**) and used to capture viral antigens which are detected via a fluorescent antibody. Each VaxArray MR assay kit (VXMR-9001, InDevR Inc.) contains two microarray slides, each with 16 replicate arrays (see **Figures 1a** and **1b)**, MR Blocking Buffer (MRBB), MR Lysis Buffer (4x), and Wash Buffers. Prior to use, VaxArray MR slides were equilibrated at 25°C for 30 min. Samples were prepared individually at 3x final concentration by lysing at 25°C for 30 min in 1x PBS + 1x MR Lysis Buffer. Each sample was further diluted in MR Blocking Buffer to 1x, and 50 μL was applied to individual arrays. Slide(s) were incubated in a humidity chamber (VX-6200, InDevR Inc.) on an orbital shaker at 65 rpm for 4 to 20 hours at 25°C. MR Blocking Buffer and antigen-specific detection label (VXMR-7634 and VXMR-7635, InDevR Inc.) was prepared and added to each array following antigen removal. Each slide was further incubated for 30 minutes in the humidity chamber at 65 rpm at 25°C before subsequent, sequential washes with Wash Buffer 1, Wash Buffer 2, 70% ethanol, and water. Slides were dried and imaged using the VaxArray Imaging System (VX-6000, InDevR Inc.). Fluorescence intensities were processed using the validated VaxArray Analysis Software utilizing the algorithms of Kuck et al. [26]. When appropriate, sample concentrations were calculated against a standard curve from the same experiment.

**Figure 1.**
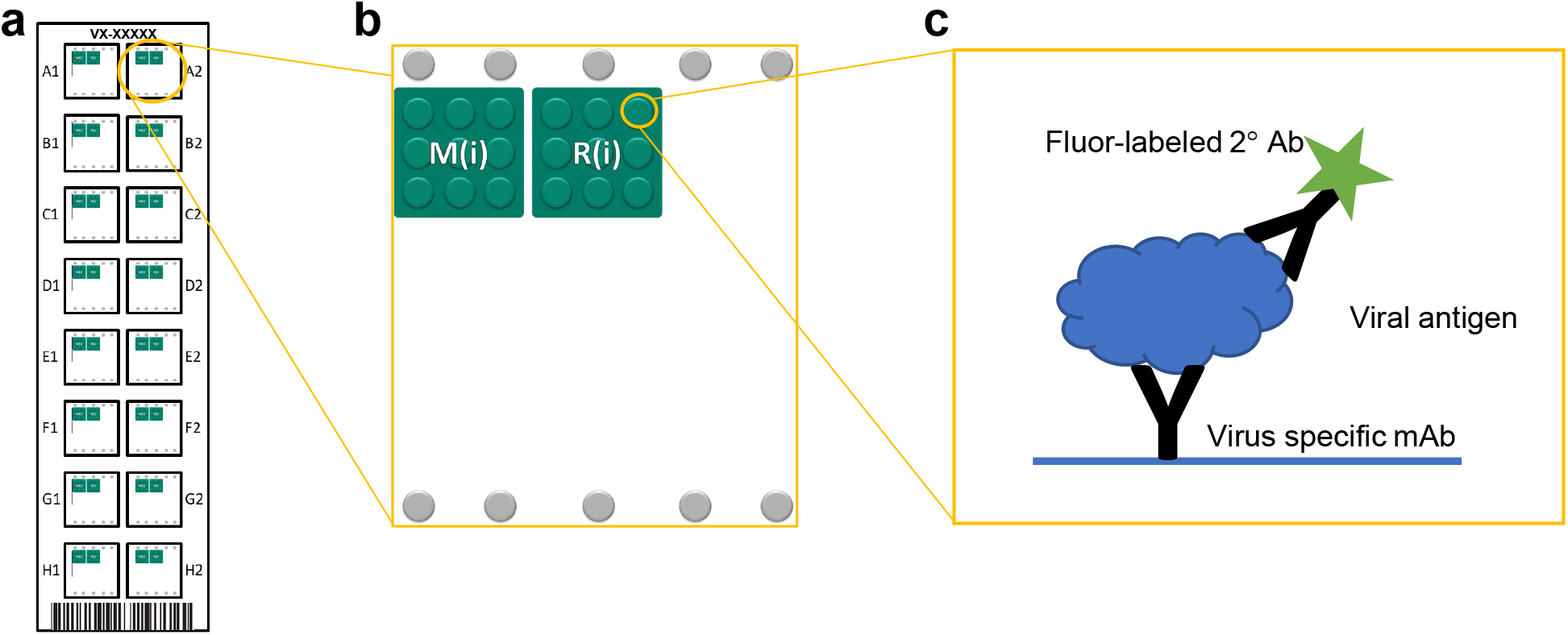
The VaxArray antibody microarray design and layout. (a) Each VaxArray slide consists of 16 identical wells, each containing the printed microarray. (b) The microarray consists of 9 spatially segregated replicate spots of each capture antibody, where M(i) denotes an anti-measles antibody and R(i) denotes an anti-rubella antibody. (c) Viral antigen, captured by the printed monoclonal antibody, is further bound by a fluorophore-conjugated secondary detection antibody, resulting in fluorescent detection and quantification.

### Cell culture

Growth medium for Vero cells (CCL-81, ATCC) consisted of Medium 199 (11150059, Gibco) supplemented to 5% fetal bovine serum (A3160401, Gibco) plus 2 mM L-Glutamine (A2916801, Gibco) and 1x Penicillin-Streptomycin (15140148, Gibco). Diluent medium for Vero cells consisted of Medium 199 (11150059, Gibco) supplemented to 2% fetal bovine serum (A3160401, Gibco) plus 2 mM L-Glutamine (A2916801, Gibco) and 1x Penicillin-Streptomycin (15140148, Gibco).

Vero cells were grown in Vero growth media in adherent cell culture flasks (Nunc EasYFlask) to 80-90% confluency. Cells were collected by washing adherent cells with sterile Dulbecco’s PBS (14190136, Gibco) followed by incubation for ~5 min with 0.25% Trypsin-EDTA (25200056, ThermoFisher). Cells were then collected, and trypsin was inactivated with an equivalent volume of supplemented growth medium.

Growth medium for Rabbit Kidney (RK13) cells (CCL-37, ATCC) consisted of Dulbecco’s Minimum Essential Medium (12-614F, Lonza) supplemented to 10% fetal bovine serum (A3160401, Gibco) plus 2 mM L-Glutamine (A2916801, Gibco) and 1x Penicillin-Streptomycin (15140148, Gibco). Diluent medium for RK13 cells consisted of Dulbecco’s Minimum Essential Medium (12-614F, Lonza) supplemented to 5% fetal bovine serum (A3160401, Gibco) plus 2 mM L-Glutamine (A2916801, Gibco) and 1x Penicillin-Streptomycin (15140148, Gibco).

Cell cultures RK13 cells were grown in RK13 growth media in adherent cell culture flasks (Nunc EasYFlask) to 80-90% confluency. Cells were collected by washing adherent cells with sterile Dulbecco’s PBS (14190136, Gibco) followed by incubation for ~5 min with 0.25% Trypsin-EDTA (25200056, ThermoFisher). Cells were then collected, and trypsin was inactivated with an equivalent volume ofsupplemented growth medium.

### Cell Culture Infectious Dose (CCID_50_)

Cell Culture Infectious Dose (CCID_50_) assays were performed at InDevR Inc. and Biological E. Ltd. following Biological E. Ltd protocols. Virus-containing samples were initially diluted in cellline matched diluent medium and further serially diluted by 0.5 log_10_ in diluent medium to create 12 virus dilutions. 50 μL diluent medium was added to all wells of treated 96-well plate(s) (161093, Nunc), and 50 μL serially diluted sample were added to each row of the plate (8 replicates each dilution). Control plates (no virus) were generated by adding an extra 50 μL diluent medium.

Vero cells were utilized for measles virus, and RK13 cells for rubella virus. Cell cultures were started ~1 week prior to executing CCID_50_. Cells were grown in a CO_2_ incubator at 37°C and 5% CO_2_ to 80-90% confluency in T175 flasks, collected by trypsin release as described above, and cell concentration determined via hemocytometer. Cells were collected, centrifuged, and reconstituted in diluent medium to create 13 mL of 1.1×10^5^ cells/mL solution per CCID_50_ plate. 100 μL cell solution was added to every well of every plate, including the control plate.

Plate(s) were covered and placed in CO_2_ incubators at 32°C (rubella) or 36°C (measles). Plates were incubated for 10 days with periodic inspection under an inverted microscope. At the end of the 10-day incubation, all wells of the plate(s) were inspected for cytopathic effects. A Spearman-Karber calculation [28,29] was applied to determine the CCID_50_ titer of each sample.

### Virus-containing samples

Three (3) lots of monovalent bulk containing live-attenuated measles CAM-70 strain, 4 lots of monovalent bulk containing live-attenuated rubella Wistar RA 27/3 strain, 3 lots of lyophilized vaccine containing measles CAM-70 and rubella Wistar RA 27/3, and 17 rubella Wistar RA 27/3-containing samples, from 2 separate lots (3 different growth conditions for one lot, and a single growth condition for the second lot), harvested at different infection times (‘harvest samples’), were obtained from Biological E. Ltd. (Hyderabad, India). A separate set of 12 harvest samples containing measles Schwarz from a single lot of material but representing 3 different growth conditions were provided by Batavia Biosciences B.V. (Leiden, Netherlands).

### Analytical sensitivity and linear dynamic range

Samples were lysed using MR Lysis Buffer (VXMR-6310, InDevR) for 30 min at 25°C in a biosafety cabinet. Samples were serially diluted in MR Blocking Buffer (VXMR-6309, InDevR) to create a 13-point standard curve. Test samples at low virus concentrations were lysed and diluted in MR Blocking Buffer prior to analysis (n=4). 50 μL of each standard was added to individual arrays on VaxArray Measles and Rubella v1.0 assay slides (VXMR-9051, InDevR), and further processed using virus-specific detection labels with slides imaged at 700 ms exposure time.

A linear regression was applied to each set of 4 adjacent standards across the 13-pt dataset with each slope and R^2^ calculated. The ULOQ was defined as the highest antigen concentration analyzed that was within a 4-point fit R^2^ > 0.95. To determine the lower limit of quantification (LLOQ), the low-concentration test samples were quantified against the matched standard curve and the average concentration and standard deviation across 4 replicates determined. LLOQ was defined as the lowest concentration with a %CV < 25% and a % difference from expected (accuracy) < 25%.

### Analytical specificity and accuracy in bivalent samples

Monovalent bulk measles CAM-70 and rubella RA 27/3 samples and bivalent mixtures of the two were prepared by serially diluting in MR Blocking Buffer, with monovalent and corresponding bivalent samples containing the same concentrations. Samples were analyzed by the VaxArray Measles and Rubella Assay as described previously. To determine specificity and accuracy of bivalent analysis, the assay response for a virus analyzed in monovalent form was compared to the same response when analyzed in the bivalent sample and the statistical difference between the two regressions compared.

### Assay precision

On three days, three separate users executed the VaxArray MR assay. On each day, a bivalent mixture of measles Schwarz (harvest sample) and rubella RA 27/3 (monovalent bulk) was prepared in PBS. The highest standard was diluted in PBS, lysed, and further diluted in MR Blocking Buffer to create the standards. In addition, 3 samples (high, middle, and low concentrations) were generated in PBS, with each lysed 8 separate times before dilution in MR Blocking Buffer. 50 μL of each standard and sample replicate were added to individual microarrays and processed as described previously. The average sample concentration and standard deviation for each sample were determined (n=8). Day-to-day results were compared for each sample, with results also normalized by their dilution factor to compare all replicates (n=72).

### Correlation and accuracy relative to CCID_50_

Monovalent harvest samples (measles Schwarz or rubella RA 27/3) were prepared and analyzed by both VaxArray CCID_50_ assay previously described. Seventeen (17) rubella harvest samples were analyzed, using a CCID_50_-characterized monovalent bulk as the VaxArray calibrant. Eleven (11) measles harvest samples were analyzed. As the difference in titer for the 11 samples was small, each sample was also further diluted twice to create 33 total samples. Measles samples were then analyzed by VaxArray and CCID_50_ using a separate harvest sample previously measured by CCID_50_ as the VaxArray calibrant.

Separately, 5 concentrated virus pools (CVP), 3 monovalent rubella RA 27/3 bulks, and 2 monovalent measles CAM-70 bulks, all from separate manufacturing lots were analyzed by VaxArray MR and CCID_50_. Additionally, 3 final vaccines (measles CAM-70 and rubella RA27/3) from separate lots were analyzed. A separate monovalent bulk for each virus from a different lot with known CCID_50_ was used as the VaxArray calibrant.

### Thermal degradation/stability indication

CAM-70 measles and RA 27/3 rubella monovalent bulk samples were aliquoted into separate glass vials, sealed with a rubber stopper and aluminum crimp-top, and frozen at −80°C. ‘Treated’ vials were then placed in a water bath at +60°C for 48 hours or in an incubator at +37°C for 24 hours, after which the ‘untreated’ vials were removed from the freezer and allowed to equilibrate to 25°C in a biosafety cabinet. Untreated and thermally treated materials were combined in specific ratios, and each prepared sample was lysed and analyzed by VaxArray. The assay response for each tested sample was compared to the fully intact sample and a % signal was determined. The percent signal relative to intact material was plotted against the % intact material and a linear regression was applied.

### Interfering substances

Separately, solutions containing 5% sucrose (S9378, Sigma), 1.125% sodium chloride (S3014, Sigma), 6.25% sorbitol (S1876, Sigma), 1% Tween 80 (BP338, Fisher Scientific), and 2% gelatin (G1393, Sigma) were prepared (w/v) in deionized water. A bivalent mixture of measles and rubella virus was prepared and combined with each vaccine-relevant matrix, resulting in samples of virus plus 0.38% sucrose, 0.90% sodium chloride, 5% sorbitol, 0.125% Tween 80, or 1.60% gelatin. A control was prepared by diluting in PBS (P3813, Sigma). Each sample was lysed using MR Lysis Buffer for 30 min, further diluted in MR Blocking Buffer to 6x LLOQ, with each added to 4 replicate arrays. A standard curve was prepared by serially diluting the PBS-control sample in MR Blocking Buffer after lysis. All slides were processed and imaged as described previously, and samples were quantified against the relevant standard curve and the average concentration of each virus in each sample determined. A student’s T-test evaluated the statistical significance of each sample relative to the control.

## RESULTS AND DISCUSSION

### Antibody pair selection

Antibodies against measles nucleoprotein and rubella E1, E2, and capsid proteins were obtained and printed on a “screening” microarray. A panel of 21 monoclonal antibodies, raised against measles (n=9) or rubella (n=12) were each printed in triplicate on the microarray Antibodies were evaluated for reactivity, specificity, limit of detection, linear dynamic range, and stability indicating capabilities, with a single antibody for each virus was down-selected for inclusion in the final assay.

### VaxArray MR assay has broad linear dynamic ranges and vaccine-relevant limits of quantification

To assess assay linearity, monovalent measles- and rubella-containing samples were analyzed to determine the ULOQ and LLOQ of the assay for each strain analyzed. As shown in **Figure 2**, the assay demonstrated good sensitivity and linearity for CAM-70 measles (**Fig 2.a**), Schwarz measles (**Fig 2.b**) and Wistar RA 27/3 rubella (**Fig 2.c**). Fig 2.c inset further highlights the linear response to lower-concentration dilutions of the rubella sample. Each strain showed linearity over ≥ 43x and resulted in LLOQ from 800 to 345 IFU/mL (**Table 1)**. Importantly, these LLOQ values are < 2,000 IFU/mL (3.3 log_10_ IFU/mL) minimum required in the final vaccine [30] and are sufficient for antigen tracking throughout vaccine manufacturing.

**Figure 2.**
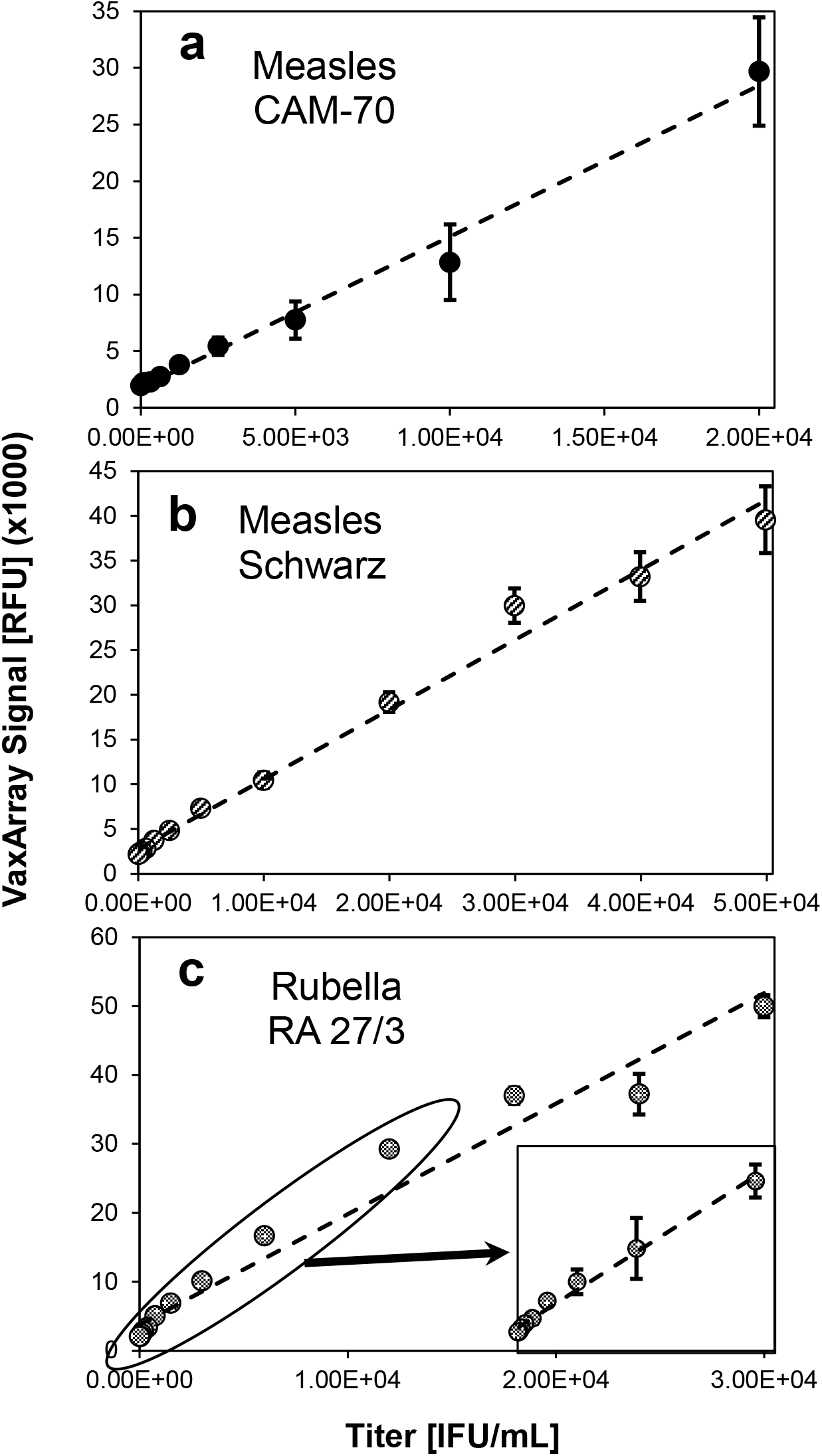
VaxArray MR assay response curves for serial dilutions of (a) measles CAM-70 strain, (b) measles Schwarz strain, and (c) rubella Wistar RA 27/3 strain. Values shown are median signal intensities, and error bars represent ± 1 standard deviation of the 9 replicate spots within the array for the relevant antibody.

**Table 1.** Limits of quantification of the VaxArray Measles and Rubella Assay

| Virus | Strain | Lower Limit of Quantification [IFU/mL] | Upper Limit of Quantification [IFU/mL] | Linear Dynamic Range |
| --- | --- | --- | --- | --- |
| Measles | CAM-70 | 800 | 100,000 | 125x |
|  | Schwarz | 345 | 50,000 | 145x |
| Rubella | Wistar RA 27/3 | 360 | 15,500 | 43x |

### VaxArray MR Assay is capable of accurate, simultaneous bivalent measles and rubella analysis

To investigate the suitability of the VaxArray MR assay for bivalent analysis, measles and rubella monovalent bulk stocks were prepared and serially diluted in monovalent form as well as combined and serially diluted as a bivalent sample (same virus concentrations present in the bivalent and corresponding monovalent samples). Each sample was analyzed and the signal responses compared to determine if the presence of the off-target virus affected accurate quantification of the target virus.

As shown in **Figure 3a**, when a measles-only sample is analyzed, only the measles-specific capture antibody shows a response while the signal intensity of the rubella-specific antibody is < 1.02x background. Similarly, as shown in **Figure 3b**, a rubella-only sample demonstrates detection only on the rubella-specific capture antibody, with the signal on the measles-specific antibody is 1.16x background. A representative image of a bivalent analysis is shown in **Figure 3c**. Direct comparison of response curves for equivalent virus concentrations in monovalent and bivalent formulations are shown in **Figures 3d** and **Figure 3e**. For measles, the linear regressions through the monovalent and bivalent data series shown in **Figure 3d** have statistically equivalent slopes and y-intercepts when considering standard error of the regressions. Specifically, a t-test showed statistical equivalence within error (p < 0.05). Similarly, the rubella-specific antibody demonstrated no difference in assay response in monovalent vs. bivalent samples **(Figure 3e)**. These results indicate that the analysis of one virus is not inhibited by and does not interfere with the presence of the other virus. The ability to simultaneously quantify both components in bivalent samples provides a distinct advantage, as the VaxArray MR assay does not require the anti-rubella neutralizing serum required for the CCID_50_ assay of measles due to the concomitant growth of rubella in Vero cells.

**Figure 3.**
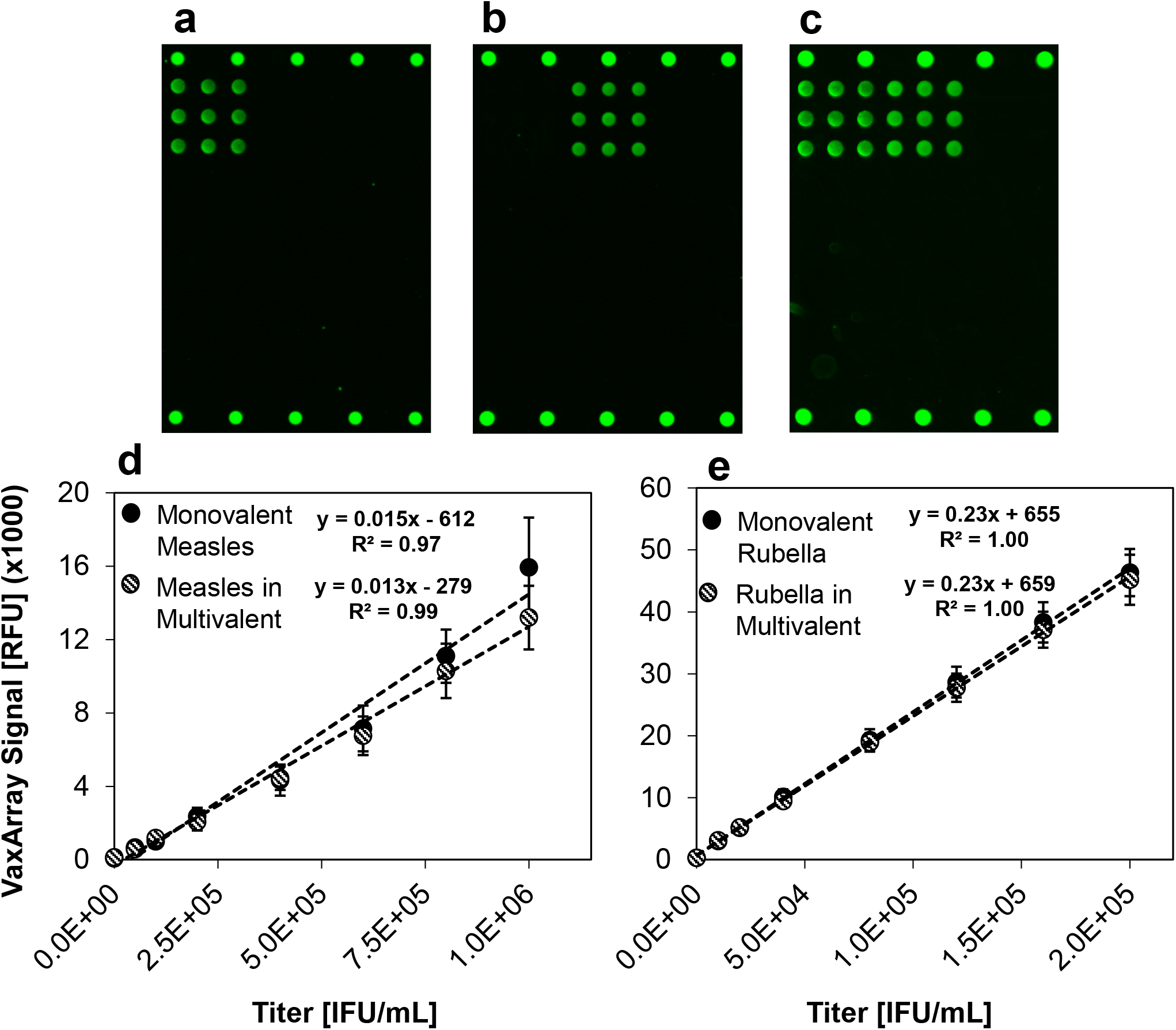
Assay response for monovalent and bivalent measles- and rubella-containing samples. In (a), an image representative of monovalent measles detection in shown. In (b), an image representative of monovalent rubella detection is shown. In (c), an image representative of bivalent measles and rubella detection is shown. In (d and e), response curves for measles and rubella in both monovalent and multivalent samples are compared, respectively. Error bars in e and f represent ± 1 standard deviation of the 9 replicate antibody spots within the array.

### VaxArray MR assay is compatible with common vaccine additives

It is also important to investigate potential interference from common stabilizers and excipients typically added to MR vaccine samples to improve shelf stability and assist in the freeze-drying process. To investigate potential interferents, a bivalent mixture of measles and rubella bulks was spiked into five matrices at vaccine-relevant concentrations[31].

Measles detection accuracy was unaffected by the presence of any of the tested additives (**Fig 4.a**), with all measurements being statistically equivalent to a PBS control. Rubella quantification was statistically unaffected for all matrices except sucrose (**Fig 4.b**). When spiked into a solution at 0.38% sucrose, rubella detection was depressed by 15% relative to the PBS control. Due to the assay’s high precision, this measurement was statistically significantly different than the PBS control (p < 0.01), however, was still within a typical 80-100% accuracy requirement, suggesting the assay produces accurate results in the presence of common MR vaccine additives. In addition, the assay’s high sensitivity allows for significant dilution of samples prior to analysis, potentially alleviating the interference by diluting the sucrose to sufficiently low levels.

**Figure 4.**
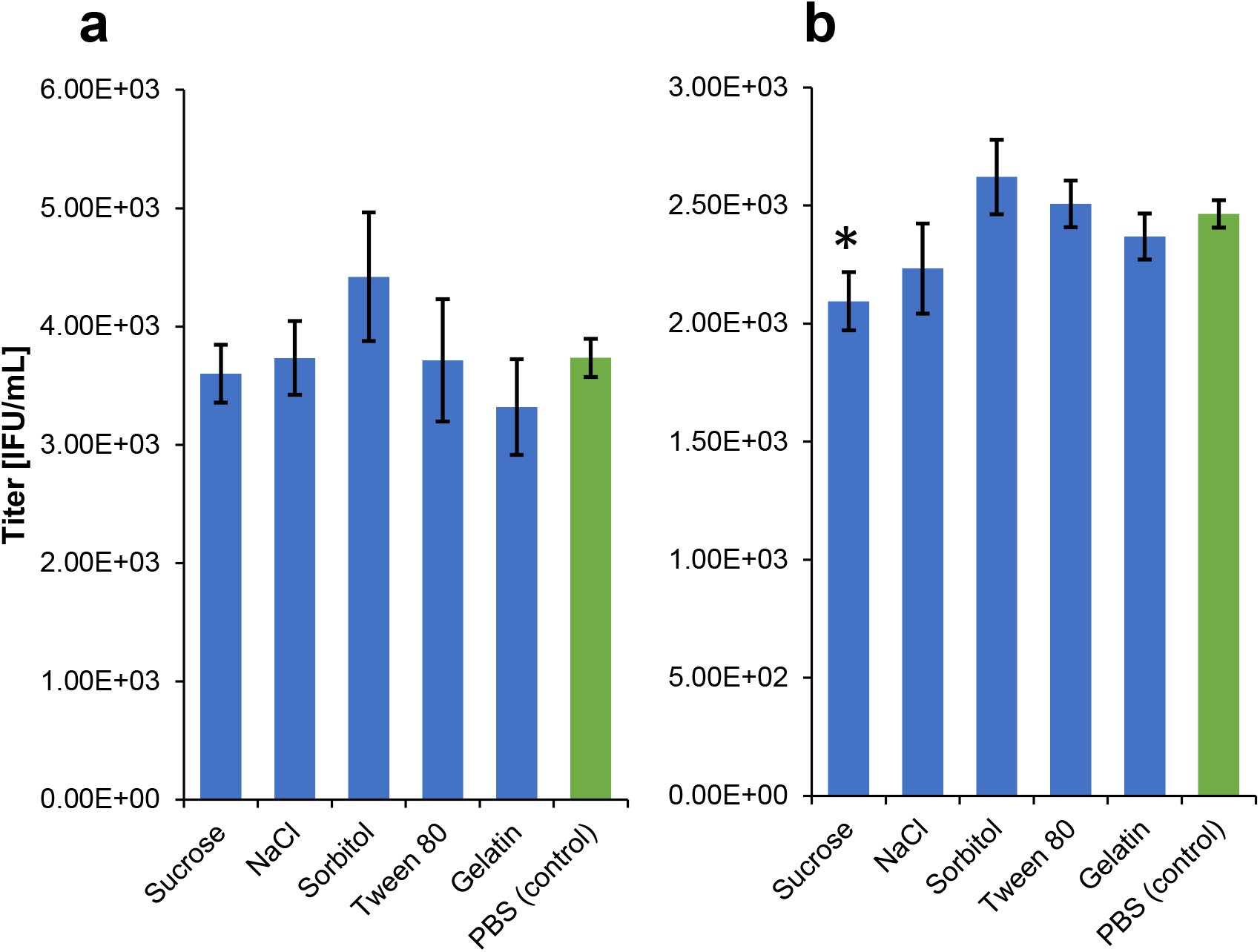
Accuracy of measles and rubella quantitation in the presence of common MR vaccine additives at vaccine relevant concentrations. Each sample was quantified against a standard curve prepared in PBS, with the average of 4 replicates of each sample reported for (a) measles and (b) rubella. Error bars represent ± 1 standard of the average. The asterisk (*) indicates a statistically significant difference from the PBS control with p < 0.01.

### VaxArray MR assay is highly reproducible

Traditional cell culture assays, such as CCID_50_, are subject to variability of up to 0.3 log_10_ IFU/mL [21–23]. This translates to relative error of up to 50%-100%. These high levels of uncertainty can cause unnecessary wastage and expensive lot rejections.

Bivalent mixtures of measles- and rubella-containing samples were analyzed at low, medium, and high concentrations relative to the LDR. Eight replicates of each bivalent sample were prepared and lysed prior to VaxArray analysis. In accordance with the ICH Guideline for Validation of Analytical Procedures [32], the study was performed by three separate users on three separate days to evaluate intermediate precision.

The assay demonstrated high precision, with %CV < 20% for all tested cases on all days (**Fig 5**). The average errors were 6.9 ± 1.5%, 9.9 ± 6.7%, and 6.5 ± 3.3% for users 1, 2, and 3, respectively, demonstrating no significant differences in the performance of the assay on different days or by different users. In addition, the error across all 72 normalized replicates was 16.2% for measles and 8.9% for rubella, demonstrating high intermediate precision and high linearity of dilution. The concentration of the ‘low’ test sample was ~6x LLOQ, indicating the assay retains high precision even at challenging concentrations. This is a significant improvement over the imprecision often experienced with CCID_50_, providing a route for same-day, precise measurements for appropriate applications.

**Figure 5.**
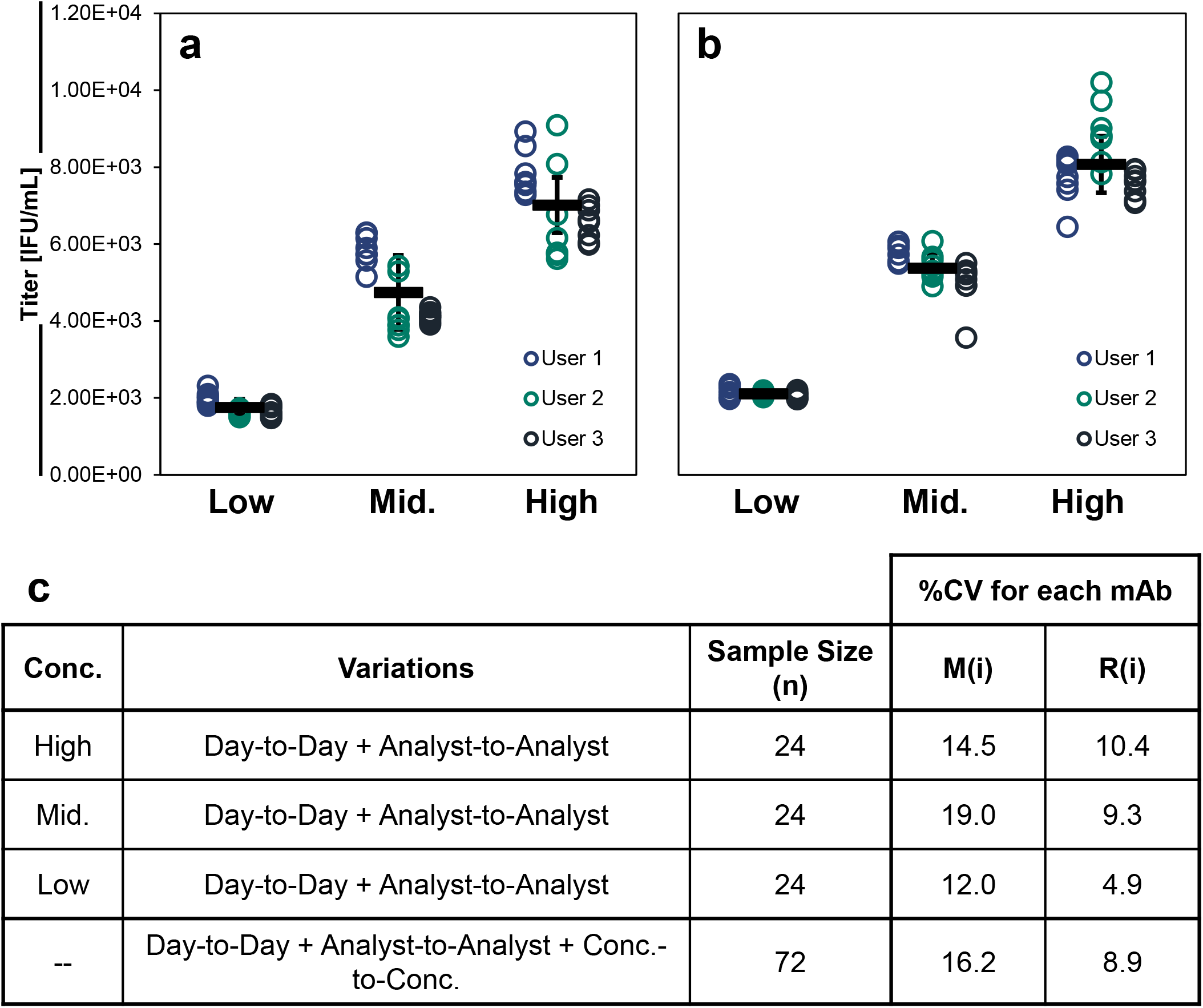
Accuracy and precision data for high, intermediate, and low concentration samples for measles (a) and rubella (b). Individual replicate results for users 1,2, and 3 are shown in blue, green, and black open circles, respectively. Solid black bars represent the overall average of the sample tested across all 3 days (n=24 replicates). Error bars represent ± 1 standard deviation of the replicates (n=24). (c) The %CV of each sample was determined across the three days/users for each sample tested for both antibodies, with all replicates normalized for dilution factor to allow a direct comparison across all 72 replicates analyzed.

### VaxArray MR assay is sensitive to antigen stability

MR vaccines are characterized by their infectivity with antigen stability an important consideration for vaccine manufacturers, as conformationally folded protein is one critical driver of infectivity, immune recognition, and vaccine efficacy. Vaccine monovalent bulks containing measles or rubella virus were subjected to 60°C for 48 hours to fully denature the viral proteins. Degraded monovalent bulk was combined with intact (non-treated) monovalent bulk at various ratios and analyzed by VaxArray MR to demonstrate the ability of the assay to measure conformationally intact protein. While antigen stability is an important consideration, retention of infectivity is also a critical consideration for live-attenuated vaccines. A second experiment in which CAM-70 measles and RA 27/3 monovalent bulk virus were treated at +37°C for 1 day was also conducted to determine the effect on the VaxArray measurement.

The VaxArray MR assay does not detect denatured proteins, as indicated bycomplete loss of signal when +60°C heat-treated virus (0% intact (undegraded) sample) was analyzed (**Fig 6, dashed circles**). As the proportion of intact sample was increased (relative to degraded sample), the assay response was linear and indicated slope of 1 and a y-intercept near 0, confirming detection of only the conformationally folded protein components in a mixture of degraded and intact sample.

**Figure 6.**
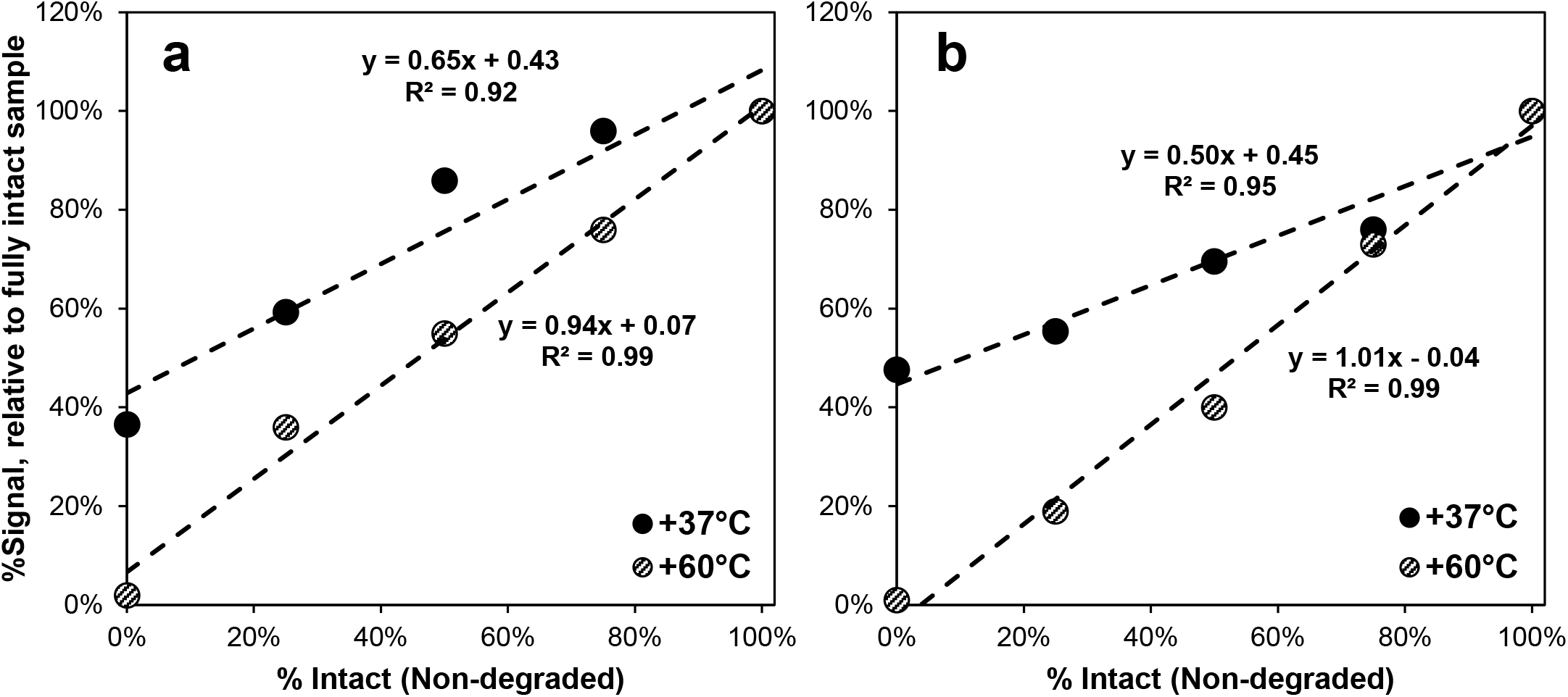
Results for monovalent measles (a) and rubella (b) bulk samples subjected to +60°C for 48 hours or +37°C for 24 hours. Mixtures with varying ratios of degraded and intact bulk were prepared and analyzed, with the signal intensity of each sample compared to that of the fully intact sample. A linear regression was applied to each set, with slopes and coefficients of regression (R^2^) as shown.

When virus was heat-treated at +37°C, the sample containing only heat-treated virus did not demonstrate complete signal loss (**Fig 6, filled circles**), but rather produced a signal of ~40% relative to the fully intact sample. As the portion of intact sample was increased (relative to degraded sample), the assay response remained linear, but the slope was < 1 for each virus, suggesting that exposing measles and rubella viruses to +37°C for 1 day does not fully denature the epitope probed by the antibodies on the microarray. In a separate study (data not shown), the same monovalent bulk virus samples incubated at +37°C for 1 day and subsequently analyzed byCCID_50_. Both bulks demonstrated complete loss of infectivity, suggesting that protein/antigen stability as measured by VaxArray is not directly predictive of infectivity for thermally degraded samples.

### VaxArray MR measurements are correlated to CCID_50_ measurements

To investigate the correlation between VaxArray and CCID_50_, crude harvest samplesfor each virus were analyzed by both assays (see Methods section for details).

VaxArray measurements were generally correlated with the CCID_50_ measurements for both viruses, with positive slopes and R^2^ of 0.81 and 0.75 for measles and rubella, respectively (**Fig 7**). However, the measured concentration is not equivalent to CCID_50_. This result is not unexpected and is likely because the VaxArray immunoassay measures conformational protein content rather than infectious dose. In the harvest samples analyzed here, there are likely levels of free protein (not associated with intact, infectious viral particles) that vary from batch-to-batch, or within a batch as a function of harvest time or growth condition. These proteins are detected by VaxArray but would not cause CPE in a CCID_50_ assay. In addition, if the ratio of free protein to infectious particles is inconsistent for all samples and the calibrant, it is expected that VaxArray measurements will not be identical to CCID_50_ measurements and therefore may be used only as a rapid measure of antigen content for these bioprocess samples. We note that the harvest samples utilized in this study were isolated from a single measles lot and 2 different rubella lots, and from numerous different manufacturing conditions within each lot. Based on preliminary studies (data not shown), analysis of samples from a single lot and a single condition (*i.e.* single standard manufacturing batch) is likely to improve the correlation.

**Figure 7.**
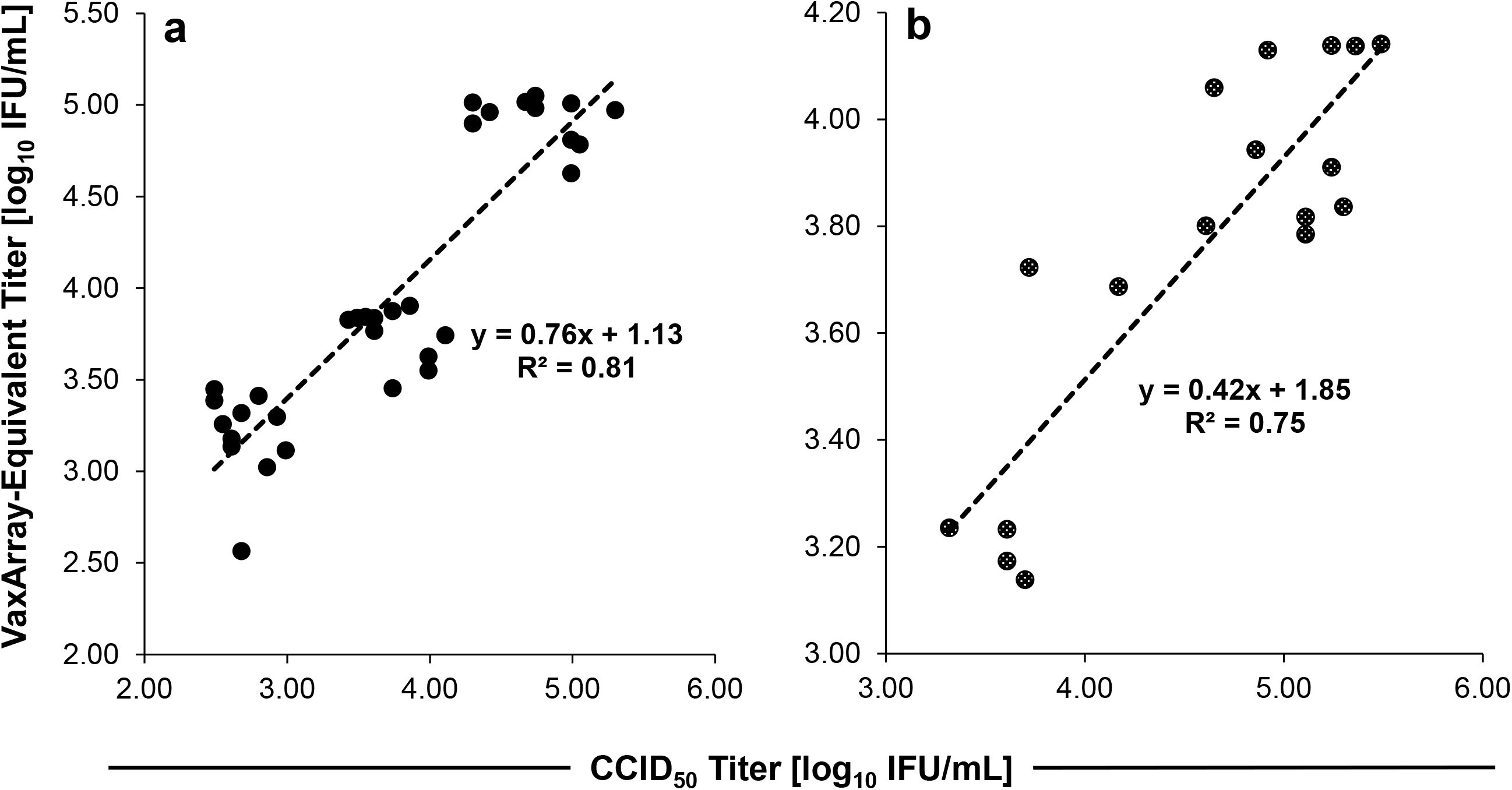
Correlations between CCID_50_ and VaxArray MR for monovalent harvest samples containing measles (a) or rubella (b), with linear regression and R^2^ shown for each dataset.

### VaxArray MR exhibits high accuracy for purified samples

To evaluate the potential use of the VaxArray MR assay with more purified downstream samples, a set of 2 monovalent measles bulks (CAM-70), and a set of 5 concentrated virus pools (CVP) and 3 monovalent rubella bulks (RA 27/3 strain, from separate manufacturing lots), were analyzed by VaxArray and CCID_50_. In addition, 3 final vaccines from separate manufacturing lots were reconstituted, and both the measles and rubella components analyzed by VaxArray and CCID_50_. Fully separate monovalent bulks from unique manufacturing lots with known CCID_50_ values were used as the VaxArray calibrant.

As summarized in **Table 2**, the VaxArray measurements for each lot of measles monovalent bulk were only 0.16 and 0.21 log_10_ IFU/mL different from the CCID_50_ measurement, within the typical CCID_50_ error of 0.30 log_10_ IFU/mL. The measles component in final vaccines, however, produced VaxArray values that were 0.36, 0.60, and 0.40 log_10_ IFU/mL different from CCID_50_ (**Table 2**), with VaxArray > infectious dose in all cases. As previously discussed, **Figure 3** indicates that it is unlikely the rubella virus is interfering with the measles measurement, as we observe identical response curves for measles in monovalent and bivalent samples. In addition, **Figure 4** indicates that the measurement accuracy is only minimally affected by the presence of any single vaccine additive. As this interference testing was done by spiking single potential interferents into monovalent liquid samples, it is possible that the multiple additives in the lyophilized vaccine affect the measles measurements differently. Therefore, a previously characterized final vaccine sample may be a more appropriate calibrant for the analysis of final vaccine samples. The measles CAM-70 analyzed is also known to exhibit aggregation, so it is possible that the virus is in different states of aggregation in the monovalent bulk as compared to the final vaccine, which may affect VaxArray differently than CCID_50_.

**Table 2.**
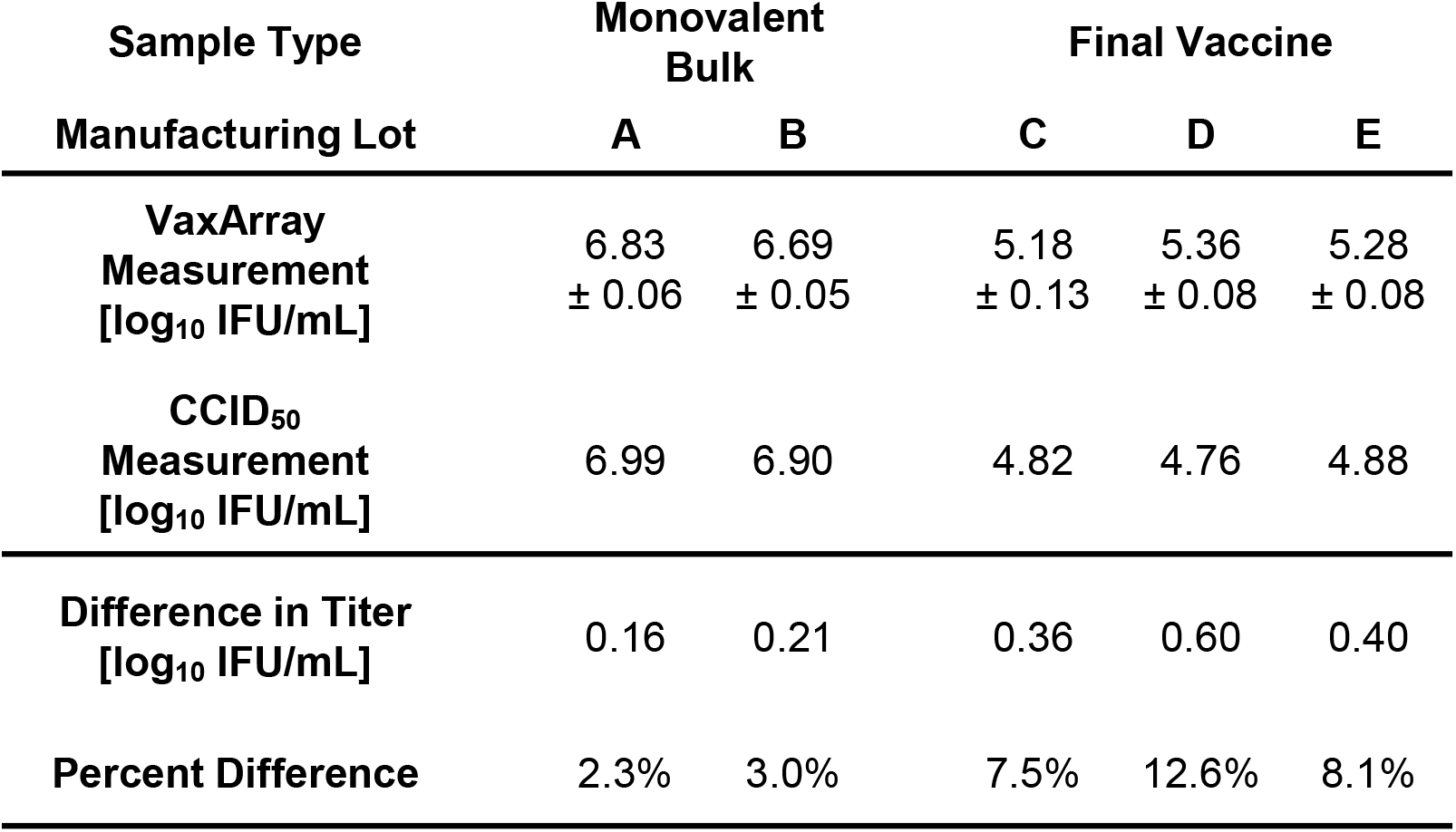
VaxArray and CCID_50_ measurements of the measles component of monovalent bulk and final vaccine samples

For rubella, the VaxArray measurements of CVP samples were within 0.05, 0.03, 0.02, 0.09, and 0.08 log_10_ IFU/mL, and the rubella monovalent bulks were within 0.16, 0.11, and 0.00 log_10_ IFU/mL of the CCID_50_ measurements (**Table 3**) for different lots of material. Rubella in final vaccines also showed good agreement with infectious dose, with VaxArray within 0.03, 0.10, and 0.16 log_10_ IFU/mL of CCID_50_ (**Table 3**), demonstrating high accuracy compared to CCID_50_ for all three types of samples. In addition, VaxArray demonstrated associated measurement errors between 0.02 to 0.18 log_10_ IFU/mL (n=2-8 replicates per sample), significantly lower than the typical 0.30 log_10_ error of the CCID_50_ assay.

**Table 3.** VaxArray and CCID_50_ measurements of the rubella component of monovalent bulk and final vaccine samples

| Sample Type | Concentrated Virus Pool (CVP) |  |  |  |  | Monovalent Bulk |  |  | Final Vaccine |  |  |
| --- | --- | --- | --- | --- | --- | --- | --- | --- | --- | --- | --- |
| Manufacturing Lot | A | B | C | D | E | B | C | F | G | H | I |
| VaxArray<br>Measurement<br>[log<sub>10</sub> IFU/mL] | 6.06<br>± 0.06 | 6.27<br>± 0.05 | 6.26<br>± 0.04 | 5.98<br>± 0.02 | 6.38<br>± 0.04 | 6.15<br>± 0.02 | 6.03<br>± 0.03 | 6.27<br>± 0.03 | 4.31<br>± 0.18 | 4.36<br>± 0.04 | 4.38<br>± 0.04 |
| CCID<sub>50</sub><br>Measurement<br>[log<sub>10</sub> IFU/mL] | 6.11 | 6.24 | 6.24 | 6.07 | 6.30 | 6.15 | 6.13 | 6.11 | 4.34 | 4.26 | 4.22 |
| Difference in Titer<br>[log<sub>10</sub> IFU/mL] | 0.05 | 0.03 | 0.02 | 0.09 | 0.08 | 0.00 | 0.11 | 0.16 | 0.03 | 0.10 | 0.16 |
| Percent Difference | 0.8% | 0.5% | 0.4% | 1.5% | 1.3% | 0.1% | 1.7% | 2.6% | 0.8% | 2.4% | 3.8% |

## CONCLUSION

While MR vaccines have proven to be extremely efficacious and generate long-lasting immunity, measles and rubella viruses still inflict severe and significant morbidity and mortality, especially in the developing world. More widespread vaccination is critical to eradicate these vaccine-preventable diseases. The current method for standardization of MR vaccine materials, including in-process harvest samples, monovalent bulks, and final vaccines, is the CCID_50_ assay which is time consuming (10-14 days), presents difficulties in the analysis of multivalent samples, and suffers from irreproducibility, creating bottlenecks and costly lot rejections that drive up vaccine cost. New tools are critical to reduce the time-to-result and increase confidence in measurements.

The VaxArray MR assay leverages the proven VaxArray technology to address these hurdles. The assay is sensitive to antigen stability and is generally correlated to infectivity for harvest samples. Quantification of antigens from one virus is unaffected by the presence of antigens from the other virus and most common MR vaccine additives, which, when combined with low limits of quantification (lower than the minimum required for vaccine samples), enable measurement of both antigens in bivalent vaccine samples.

In conjunction with a pre-calibrated internal standard, the VaxArray MR assay demonstrated high accuracy relative to CCID_50_ for purified samples including concentrated virus pools and monovalent bulks but lower accuracy with crude harvest samples. However, these crude samples spanned multiple manufacturing lots and often different growth conditions within a lot. Further analysis of a larger number of harvest samples from a single lot and single growth condition is warranted to determine if the correlation is improved. Quantification in monovalent bulks using a previously characterized monovalent bulk as the calibrant yields more accurate measurements than in a crude harvest sample, in part, because the matrix is matched. Furthermore, after crude harvest material undergoes purification and concentration steps to arrive at monovalent bulk, the ratio of infectious virions to non-infectious material is likely more consistent. And while VaxArray MR assay results for harvest samples are correlated but not equivalent to CCID_50_ measurements, when paired with an internal standard previously calibrated by CCID_50_, the VaxArray MR assay may still provide utility in selecting harvests for pooling prior to purification in a fraction of the time relative to CCID_50_.

In addition to a short time-to-result (5 hours vs 2 weeks), VaxArray MR exhibited improved reproducibility and precision with overall %CV of 15%. This ~10-fold improvement over CCID_50_ could alleviate a major bottleneck in the characterization and quantification of MR vaccine samples by reducing the incidence of costly lot rejections. Further work to develop a sample pre-treatment protocol to improve the correlation with CCID_50_ harvest samples and for vaccine stability investigations would further increase the utility of the assay throughout the MR vaccine manufacturing process. While the focus of this work is on measles and rubella (MR) vaccines, future addition of mumps and varicella-zoster to the assay would broaden applicability.

## DATA AVAILABILITY STATEMENT

All relevant data from this study are available from the authors.

## ACKNOWLEDGEMENTS

This work was supported, in whole or in part, by the Bill & Melinda Gates Foundation [INV-004629]. Under the grant conditions of the Foundation, a Creative Commons Attribution 4.0 Generic License has already been assigned to the Author Accepted Manuscript version that might arise from this submission. We also thank Batavia Biosciences B.V. for providing measles Schwarz harvest samples for analysis under a material transfer agreement.

## DECLARATION OF INTERESTS

K. Rowlen and E. Dawson are InDevR Inc. stockholders. Biological E. Ltd. was provided a VaxArray Imaging System and kits used for analysis at no charge. All other authors are current or former InDevR Inc. employees or employees of Biological E. Ltd. but have no conflicts of interest.

## AUTHOR CONTRIBUTIONS

Mr. Gillis, Dr. Byrne-Nash, Dr. Dawson, and Ms. Taylor were responsible for experimental design, data interpretation. Mr. Gillis prepared the manuscript; all authors provided edits. Mr. Gillis, Mr. Miller, Ms. Thomas, and Mr. Panchakshari executed experiments. Dr. Rowlen, Dr. Byrne-Nash, and Mr. Gillis are inventors on related intellectual property. Dr. Rowlen and Dr. Dawson provided scientific guidance and program oversight. Dr. Dawson, Dr. Rowlen, and Dr. Senthilkumar Manoharan provided technical support and manuscript review.

